# The Cancer Testes Antigen, HORMAD1, is a Tumor-Specific Replication Fork Protection Factor

**DOI:** 10.1101/2023.01.31.526348

**Authors:** Luis Reza Herrera, Kathleen McGlynn, Zane A. Gibbs, Anthony J. Davis, Angelique W. Whitehurst

## Abstract

Tumors frequently activate the expression of genes that are only otherwise required for meiosis. HORMAD1, which is essential for meiotic recombination in multiple species, is expressed in over 50% of human lung adenocarcinoma cells (LUAD). We previously found that HORMAD1 promotes DNA double strand break (DSB) repair in LUAD. Here, we report that HORMAD1 takes on an additional role in protecting genomic integrity. Specifically, we find HORMAD1 is critical for protecting stalled DNA replication forks in LUAD. Loss of HORMAD1 leads to nascent DNA degradation, an event which is mediated by the MRE11-DNA2-BLM pathway. Moreover, following exogenous induction of DNA replication stress, HORMAD1 deleted cells accumulate single stranded DNA (ssDNA). We find that these phenotypes are the result of a lack of RAD51 and BRCA2 loading onto stalled replication forks. Ultimately, loss of HORMAD1 leads to increased DSBs and chromosomal aberrations in response to replication stress. Collectively, our data support a model where HORMAD1 expression is selected to mitigate DNA replication stress, which would otherwise induce deleterious genomic instability.

## INTRODUCTION

Cancer Testes Antigens (CTAs) are a collection of ~250 proteins, which are defined by an expression pattern that is normally restricted to reproductive tissues, but also abnormally activated in a wide variety of tumors (Almeida et al., 2009; Gibbs and Whitehurst, 2018; Simpson et al., 2005). These proteins are referred to as “antigens” because testes are an immune privileged site and peptide antigens derived from these proteins can evoke an immune response when presented on the surface of somatic cells (Atanackovic et al., 2006; Fijak and Meinhardt, 2006; van der Bruggen et al., 1991). Thus, CTAs have long been touted as ideal immunotherapeutic targets; an idea that is supported by the success of infusion of patient derived, ex vivo expanded T-cells targeting CTAs (D’Angelo et al., 2018; Hoyos et al., 2022; Hunder et al., 2008).

Recent studies suggest that CTA proteins are not innocuous when expressed in tumor cells. Instead, a number of reports have identified individual CTAs that are directly engaged in support of neoplastic behaviors, including: degradation of tumor suppressors, promoting resistance to apoptosis, enhancing oxidative phosphorylation, and reprogramming transcriptional networks (Cappell et al., 2012; Cheng et al., 2020; Epping et al., 2005; Gallegos et al., 2019; Gibbs et al., 2020; Gibbs and Whitehurst, 2018; Maxfield et al., 2015; Michael et al., 2015; Whitehurst et al., 2010). The revelation that these anomalously expressed testes proteins are required for oncogenic behaviors establishes a previously unappreciated aspect of the tumor cell regulatory environment. Moreover, these proteins represent direct intervention targets, which based on their biased expression pattern may have a broad therapeutic window.

Within the CTA family, there are seven proteins with highly conserved functions in meiosis: HORMAD1, HORMAD2, SPO11, SYCE1, SYCE2, SYCP1, SYCP2 and SYCP3 (Baudat et al., 2000; Bolcun-Filas et al., 2007; Bolcun-Filas et al., 2009; de Vries et al., 2005; Keeney et al., 1997; Kogo et al., 2012; Shin et al., 2010; Yang et al., 2006; Yuan et al., 2000). In mammals, these proteins are essential for homologous recombination (HR), which occurs during meiotic prophase 1. Specifically in sperm, HORMAD1 and HORMAD2 are recruited to chromosomes, a process required for the generation of DNA double strand breaks (DSBs) catalyzed by the SPO11 endonuclease (Daniel et al., 2011; Keeney *et al.*, 1997; Kogo *et al.*, 2012; Panizza et al., 2011; Schramm et al., 2011; Shin *et al.*, 2010). These breaks, after single strand processing, are the basis for homology search and alignment of chromosomal homologues. Immediately following alignment, the CTAs SYCE1, SYCE2, SYCP1, SYCP2 and SYCP3 help form a proteinaceous bridge between homologous chromosomes termed the synaptonemal complex (SC)(Bolcun-Filas *et al.*, 2007; de Vries *et al.*, 2005; Li Yuan, 2000; Yang *et al.*, 2006). The SC mediates DSB repair and the formation of crossovers that ultimately permit exchange of maternal and paternal genetic information. Inability to undergo meiotic recombination prevents germ cell maturation. Indeed, deletion of any one of these CTAs in mice leads to infertility in both sexes (Baudat *et al.*, 2000; Bolcun-Filas *et al.*, 2007; Bolcun-Filas *et al.*, 2009; de Vries *et al.*, 2005; Geisinger and Benavente, 2016; Kogo *et al.*, 2012; Li Yuan, 2000; Romanienko and Camerini-Otero, 2000; Shin *et al.*, 2010; Yang *et al.*, 2006; Yuan *et al.*, 2000). Notably, these knockout mice develop normally without additional detectable defects, indicating the specificity of these CTAs to meiotic processes.

Expression of each of these “meiotic” CTAs has been demonstrated in a wide variety of tumors and immune reactivity is present in a subset of tumors (Atanackovic *et al.*, 2006; Chen et al., 2005a; Chen et al., 2005b; Liu et al., 2012; Neumann et al., 2005; Niemeyer et al., 2003; Taguchi et al., 2014; Tureci et al., 1998). Given their highly specialized roles in meiosis, it is difficult to predict ab initio their function in cancer cells. However, a few details have begun to emerge indicating that these proteins regulate genomic integrity positively and/or negatively in tumors. Specifically, we and others have found that HORMAD1 expression correlates with increased mutation burden and chromosomal scarring in tumors (Gao et al., 2018; Liu et al., 2020a; Nichols et al., 2018; Tarantino et al., 2022; Watkins et al., 2015; Xian Wang, 2018; Zong et al., 2021). In addition, HORMAD1 loss in cancer cells can increase sensitivity to DNA damaging agents including radiation, camptothecin, and cisplatin (Gao *et al.*, 2018; Liu *et al.*, 2020a; Nichols *et al.*, 2018; Watkins *et al.*, 2015). Importantly, the loss of HORMAD1 prevents growth of tumors in vivo (Nichols *et al.*, 2018). Collectively, reports on HORMAD1 function indicate it can impact multiple DNA repair pathways in a context specific manner (Gao *et al.*, 2018; Liu *et al.*, 2020a; Nichols *et al.*, 2018; Tarantino *et al.*, 2022; Watkins *et al.*, 2015; Xian Wang, 2018; Zong *et al.*, 2021). Whether HORMAD1 promotes and/or prevents genomic instability remains unclear.

Here, we report that HORMAD1 can directly impact tolerance of DNA replication stress. We used tandem mass spectrometry to identify HORMAD1 interacting partners in lung adenocarcinoma (LUAD) cells. We identified that HORMAD1 associates with multiple proteins that mediate DNA replication and the response to replication stress, which drove us to examine if HORMAD1 functions in these processes. Indeed, HORMAD1 is crucial for the protection of stalled DNA replication forks, as its loss leads to nascent DNA degradation in multiple LUAD cell lines. We find that HORMAD1 is essential for the recruitment of the fork protection factors, RAD51 and BRCA2, to reversed forks. Loss of HORMAD1 leads to nucleolytic-mediated degradation mediated by the MRE11-DNA2-BLM pathway. In the absence of HORMAD1, replication forks undergo excessive DNA degradation and the accumulation of chromosomal aberrations. Our findings indicate that HORMAD1 can promote genomic integrity through the protection of nascent DNA during replication stress, which would otherwise lead to degradation and genomic instability.

## RESULTS

### HORMAD1 protects stalled replication forks from degradation

Depletion of HORMAD1 in LUAD cells increases sensitivity to DNA damaging agents, suggesting a function in the cellular response to genotoxic stress (Gao *et al.*, 2018; Nichols *et al.*, 2018; Watkins *et al.*, 2015). To further elucidate the molecular mechanisms of HORMAD1 in tumor cells, we performed HORMAD1 immunoprecipitations followed by nano-HPLC mass spectrometry to identify endogenous HORMAD1 complexes in two LUAD cell lines that express robust amounts of HORMAD1: H1395 and A549. This approach returned a cohort of proteins with annotated functions in DNA replication and mediators of replication stress response. Specifically, HORMAD1 forms complexes with the DNA replication factors PCNA, RFC3, RFC4, ATAD5, and multiple MCM family members; as well as the replication stress response factors WRNIP1 and PARP1 (Supplemental Table 1)(Leuzzi et al., 2016; Park et al., 2019; Thakar et al., 2020).

The interaction data implicates HORMAD1 in DNA-replication-associated processes in LUAD cells. To test this possibility, we generated HORMAD1 knockout H1395 cells via CRISPR/Cas9. HORMAD1 loss was verified by immunoblotting (Figure S1A). Knockout of HORMAD1 did not affect the cell cycle distribution or growth rate (Figures S1A and B). To assess HORMAD1’s potential contribution to DNA replication-associated activities, we employed DNA fiber assays (Quinet et al., 2017). First, we determined if HORMAD1 regulates DNA replication in unperturbed conditions by sequentially pulse-labelling cells with the thymidine analogues 5-iodo-2’-deoxyuridine (IdU) and 5-chloro-2’-deoxyuridine (CldU). Comparison of the length of red and green tracks did not reveal any differences between sgCTRL and sgHORMAD1, indicating that baseline DNA replication-associated fork progression was not impacted by the loss of HORMAD1 (Median fiber length sgCTRL= 19.50μm, Median fiber length= 19.86μm) (Figure 1A). Next, we examined if HORMAD1 affected replication fork behavior after the induction of replication stress via treatment with hydroxyurea (HU). Specifically, the cells were pulse-labeled with IdU, then treated with HU, and finally pulse-labeled with CldU. DNA fiber analysis showed that exposure to HU induced a similar number of stalled forks in the sgCTRL and sgHORMAD1 cells (Figure 1B, i). Furthermore, sgCTRL and sgHORMAD1 cells exhibited a similar percentage of restarted forks (~79.7% vs 80.6%), suggesting that HORMAD1 is not required for resumption of DNA replication following stress (Figure 1B, ii). However, closer examination of the DNA fibers revealed that the length of the IdU labeling was significantly shorter in the sgHORMAD1 cells compared to the sgCTRL, suggesting that HORMAD1 is required for the integrity of the fork during replication stress (Figure 1B, iii). To assess if HORMAD1 protects stalled forks specifically, we monitored the stability of nascent replication strands. The DNA was labelled sequentially with IdU then CldU followed by HU. In this scenario, the length of the CldU track reflects stability of newly synthesized DNA; therefore, the ratio of the length of CldU to IdU tracks were measured. In non-HU exposed samples, sgCTRL and sgHORMAD1 cells exhibited a similar ratio (sgCTRL CldU/IdU ratio: 0.75 vs sgHORMAD1 CldU/IdU ratio: 0.76). Strikingly, after exposure to HU, we observed a substantial decrease in the CldU/IdU ratio in the sgHORMAD1 cells, reflecting a shortening of the CldU track (sgCTRL CldU/IdU ratio:0.75 vs sgHORMAD1 CldU/IdU ratio: 0.50) (Figure 1C). This result indicates that HORMAD1 promotes the protection of nascent DNA at stalled replication forks from degradation. This defect is specific to HORMAD1, as ectopic expression of HORMAD1 was sufficient to rescue the decreased ratio (sgHORMAD1 CldU/IdU ratio: 0.64 vs HORMAD1-V5 CldU/IdU ratio: 0.86) (Figure 1D). This defect was also recapitulated when HORMAD1 was depleted with siRNAs (Figure 1E, left panel). To determine if this phenomenon is conserved in multiple genetic backgrounds, we depleted HORMAD1 in seven additional LUAD cell lines. We found that HORMAD1 was required for fork protection in five of the eight cell lines tested (Figures 1E and S2C). Collectively, these findings indicate that HORMAD1 protects HU-induced stalled replication forks.

**Figure 1.**
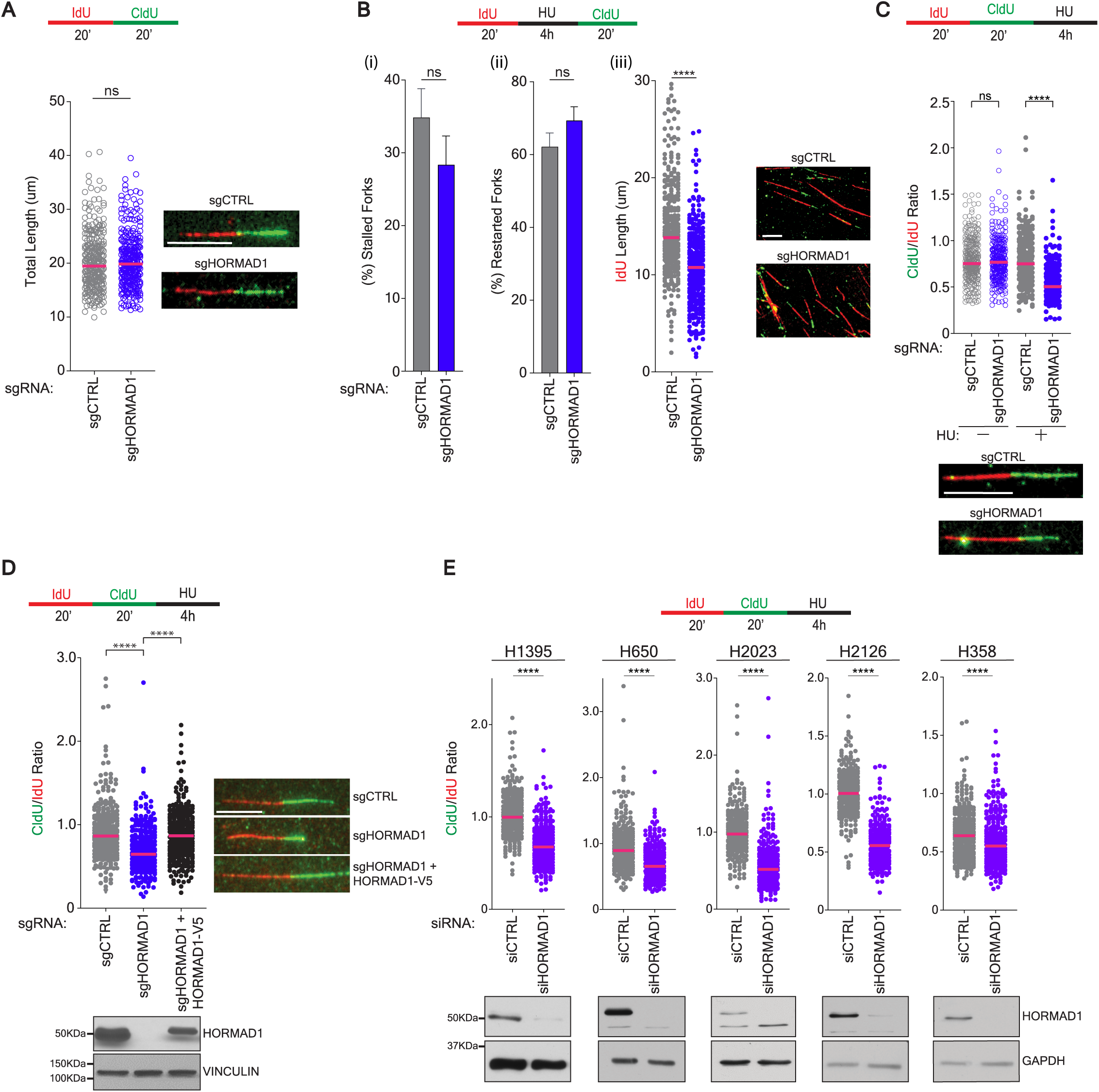
HORMAD1 protects stalled replication forks from degradation. (**A**) A DNA fiber assay was performed in indicated H1395 cells according to the schematic (top). Left: Each circle represents the length of individual tracts with median indicated. *P* value calculated by Mann-Whitney test; n≥3. Right: Representative image of a single DNA fiber. Scale bar = 10μm (**B**) A DNA fiber assay was performed in indicated H1395 cells according to the schematic (top). (i) Bars represent the mean of forks with only IdU labelling (error bars represent Standard Error of the Mean (SEM)) as a percentage of all forks observed. *P* value calculated by an unpaired t-test; n≥3 (ii): Bars represent the mean (error bars represent SEM) of tracts labeled with both IdU & CldU as a percentage of total tracts observed. *P* value calculated by an unpaired t-test; n≥3. (iii): Graph represents the IdU length of individual fibers. Median is indicated. *P* value calculated by Mann-Whitney test; n≥3. Right: Representative images of fibers are shown. Scale bar = 10μm. (**C**) Indicated H1395 cells were labelled and treated as indicated in schematic (top) followed by a DNA fiber assay. Each dot represents the ratio of CldU length to IdU length for an individual tract. Median is indicated. *P* value was calculated by Mann-Whitney test; n ≥3. Below: Representative images of fibers are shown, Scale bar = 10μm. (**D**) As in (C). Left: Graph represents the CldU/IdU ratio of individual DNA strands with the median indicated. *P* value calculated by Mann-Whitney test n≥3. Right: Representative images of fibers are shown. Scale bar 10μm. Below: Parallel lysates were obtained, and a western blot performed for indicated antibodies. (**E**) HORMAD1 was depleted for 72 hours is indicated LUAD cell lines. Cells were labelled and treated according to the schematic (top). Graphs represent the median CldU/IdU ratio for individual tracts. Median is indicated. *P* value calculated by Mann-Whitney test; n≥3. Below: Parallel lysates were obtained, and a western blot performed for indicated antibodies.

### HORMAD1 promotes recruitment of RAD51 and BRCA2 to stalled replication forks

The response to replication stress is complex and includes checkpoint activation, stalled fork stabilization, fork remodeling, fork reversal, and restart of replication. Our data suggest that HORMAD1 promotes protection of stalled replication forks. Thus, we first assessed replication stress induced checkpoint activation in the absence of HORMAD1. Here HU-induced phosphorylation of ATR (p-Thr1989) and CHK1 (p-Ser345) were unchanged in sgHORMAD1 cells as compared to sgCTRL indicating that HORMAD1 acts downstream of the signaling events induced by replication stress (Figure S2A).

Following checkpoint activation, stalled replication forks are stabilized through a fork reversal mechanism where nascent DNA hybridizes (Berti et al., 2020). Stability of this reversed fork structure is dependent upon PARP1 activity. Indeed, when H1395 cells were pre-treated with the PARP1 inhibitor, Olaparib, the fork degradation phenotype in cells lacking HORMAD1 was rescued (Figure 2A). This result implicates HORMAD1 in protecting reversed forks during replication fork reversal due to slowing or stalling of the fork. Previous studies indicate that fork reversal is mediated by different translocases and/or helicases, including SMARCAL1, ZRANB3, HLTF, and FBH1 (Liu et al., 2020b). To assess which translocase/helicase is responsible for generating the reversed fork that HORMAD1 helps protect, we individually depleted SMARCAL1, ZRANB3, HLTF, ZRANB3, and FBH1 in siCTRL and siHORMAD1 cells and measured DNA degradation. Only depletion of FBH1 rescued fork degradation in the siHORMAD1 cells, suggesting a dependency on HORMAD1 for in protection of FBH1 reversed forks (Figure 2B, S2B).

**Figure 2.**
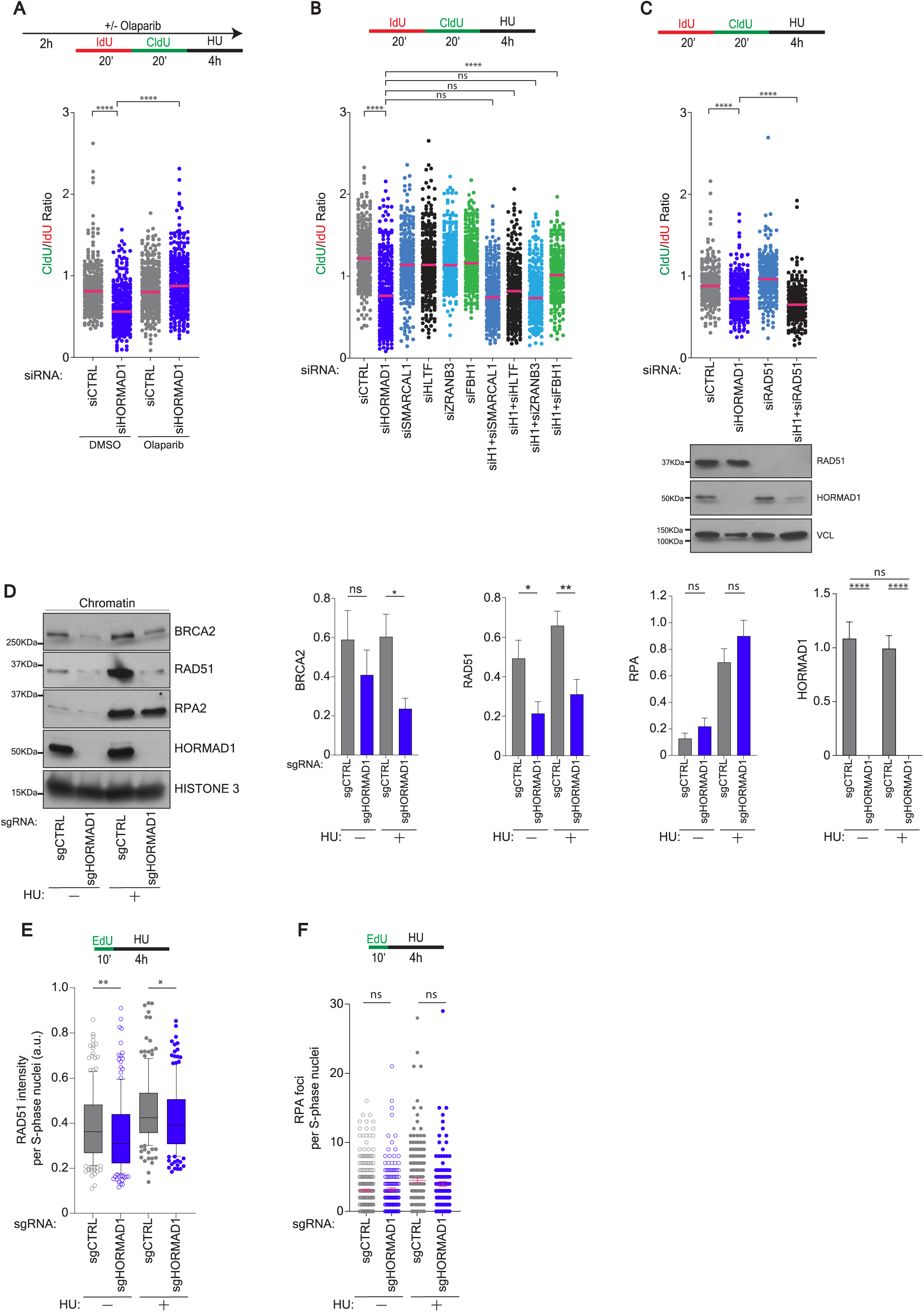
HORMAD1 promotes recruitment of RAD51 and BRCA2 to stalled replication forks. **(A)** H1395 cells were transfected with indicated siRNAs for 72 hours. Cells were treated according to the schematic (top) followed by a DNA fiber assay as in 1C. Median indicated. *P* value calculated by Mann-Whiney, n≥3. **(B)** H1395 cells were transfected with indicated siRNAs for 72 hours prior to treatment (top) and DNA Fiber assay as in Figure 1C. Median is indicated. *P* value calculated by Mann Whitney t-test; n≥3. **(C)** Top: As in (B). Below: Parallel whole cell lysates were obtained and western blotted with indicated antibodies. **(D)** Left: Chromatin fractions of indicated H1395 cells exposed to PBS or HU were western blotted for indicated antibodies. Right: Graphs represent the mean band intensity. Error bars represent SEM. *P* value calculated by Mann-Whitney, except for BRCA2 and RAD51 which was unpaired t-test; n≥3. **(E)** Indicated H1395 cells were labelled with EdU then exposed to HU or processed for IF with anti-RAD51 and anti-EdU. Box plot was derived by measuring the mean RAD51 nuclear intensity in EdU positive cells. *P* value calculated by Mann-Whitney test; n≥3. **(F)** As in **(D)** except samples immunostained for RPA. Graph represents the mean number of foci per nucleus. Mean SEM is indicated. *P* value calculated by Mann-Whitney test; n≥3.

Next, we investigated the involvement of RAD51 in this process, as it both promotes fork reversal and protects the regressed fork from nucleolytic cleavage. To our surprise, depletion of RAD51 did not rescue nascent strand degradation in the HORMAD1-deficient cells (Figure 2C). This result drove us to postulate that HORMAD1 protects stalled replication forks by promoting the recruitment to and/or stabilization of RAD51 at the reversed fork. To assess this, we examined accumulation of RAD51, BRCA2, and RPA to the chromatin fraction following HU treatment. Immunoblotting of isolated chromatin fractions with and without HU treatment revealed that HORMAD1 is constitutively bound to chromatin and does not undergo an appreciable change during replication stress (Figure 2D). In sgHORMAD1 cells, we observed a substantial decrease in recruitment of RAD51 and BRCA2 to chromatin as compared to sgCTRL cells at baseline and more dramatically following HU treatment (Figure 2D). RPA abundance was unchanged under all conditions (Figure 2D). Single cell analysis recapitulated these findings, showing that RAD51, but not RPA, signal is significantly decreased in sgHORMAD1 cells in S-Phase nuclei compared to sgCTRL cells (Figure 2E and F). Together, these findings indicate that HORMAD1 protects the nascent DNA of FBH1-mediated reversed forks by promoting RAD51 and BRCA2 chromatin localization.

### HORMAD1 protects the replication fork from nascent DNA degradation by the MRE11-DNA2-BLM enzymatic axis

Numerous studies have reported that MRE11 activity is responsible for degrading HU-induced stalled forks in BRCA2-defective cells (Datta et al., 2021; Lemacon et al., 2017; Schlacher et al., 2011). Since HORMAD1-deficient cells exhibited degradation similar to that observed in BRCA2-defective cells, we examined if MRE11 nuclease activity is responsible for fork degradation in cells lacking HORMAD1. Here, we co-depleted HORMAD1 and MRE11 and measured degradation (Figure 3A). Indeed, MRE11 depletion fully rescued degradation upon HORMAD1 depletion and HU exposure (siHORMAD1 median: 0.64 vs siHORMAD1+ siMRE11 median= 0.98). As MRE11 exhibits both endo- and exonuclease, we incubated siCTRL and siHORMAD1 cells with Mirin (exonuclease inhibitor) or PFM01 (endonuclease inhibitor) prior to IdU-CldU labeling and HU exposure to identify which activity is responsible for fork degradation. We observed that pretreatment with PFM01, but not Mirin, significantly rescued the fork degradation (Figure 3B). These data illustrate that HORMAD1 protects stalled forks from endonucleolytic attack by MRE11.

**Figure 3.**
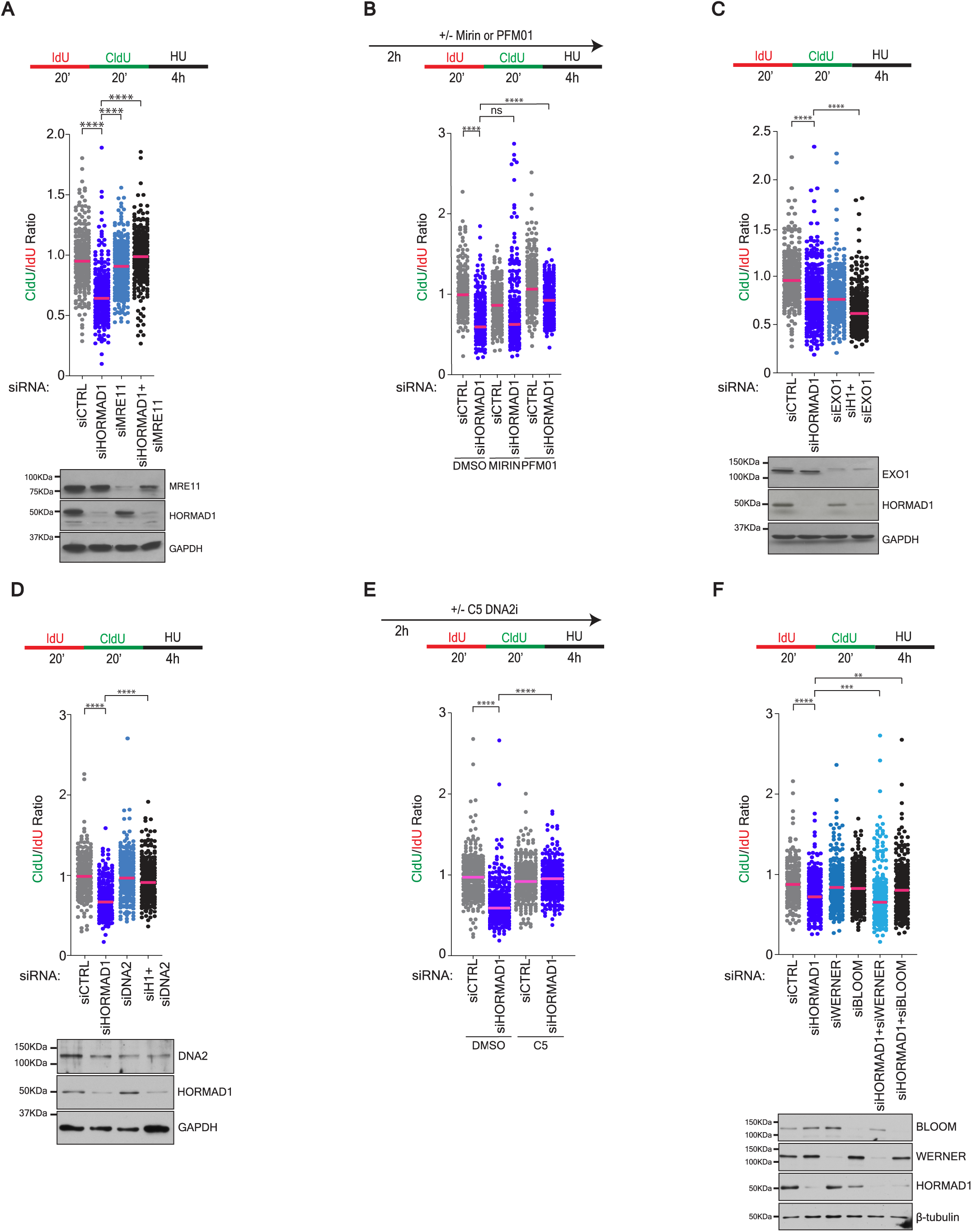
HORMAD1 protects the replication fork from nascent DNA degradation by the MRE11-DNA2-BLM enzymatic axis. (**A**) H1395 cells were transfected with indicated siRNAs for 72 hours before labelling according to the schematic (top) and a DNA fiber assay. Graph represents the CldU/IdU ratio for DNA fibers with the median indicated. *P* value calculated by Mann-Whitney t-test; n≥3. Below: Parallel lysates were obtained and western blotted with indicated antibodies. (**B**) H1395 cells were transfected with indicated siRNAs for 72 hours. Cells were exposed to vehicle or inhibitors for two hours prior to labelling according to schematic (top). Graph as in **(A)**. *P* value calculated by Mann Whitney t-test; n≥3. (**C & D**) As in A. (**E**) H1395 cells were transfected with indicated siRNAs for 72 hours. Cells were exposed to DMSO or DNA2i C5 for 2 hours prior to labelling (top) and DNA fiber assay. Graph represents the CldU/IdU ratio for individual fibers with median indicated. *P* value calculated by Mann Whitney t-test; n≥3. (**F**) As in A.

MRE11 endonuclease activity often acts in concert with the long-range nucleases DNA2 and EXO1 (Cejka and Symington, 2021). Thus, we co-depleted HORMAD1 and either EXO1 or DNA2 and measured DNA degradation. This analysis revealed HORMAD1 protects specifically against DNA2 function, but not EXO1 (Figures 3C and D). We observed comparable results using the DNA2 exonuclease inhibitor, C5 (Figure 3E). DNA2 requires the Bloom (BLM) or Werner (WRN) helicase for unwinding DNA prior to digestion (Sturzenegger et al., 2014; Thangavel et al., 2015). We depleted each helicase in combination with HORMAD1. In this setting, BLM, but not WRN, is sufficient to rescue the nascent DNA degradation observed upon HORMAD1 depletion (Figure 3F). Collectively, these findings indicate that HORMAD1 protects FBH1-reversed replication forks from MRE11-DNA2-BLM nuclease attack by promoting the recruitment of the fork protection factors RAD51 and BRCA2.

### HORMAD1 mitigates DNA damage during DNA replication stress

Given our findings that HORMAD1 protects DNA from degradation, we hypothesized that HORMAD1 may promote genomic integrity. To evaluate this possibility, we first measured γH2AX signal (an indirect marker for DSBs) in cells lacking HORMAD1. We observed an increase in signal in sgHORMAD1 cells compared to sgCTRL at baseline, which supports the notion that HORMAD1 protects against replication stress-associated DNA damage, which is likely continuously occurring in tumor cells (Figure 4A). In response to HU the γH2AX signal was markedly increased in the sgHORMAD1 cells compared to sgCTRL (Figure 4A). Furthermore, in the alkaline comet assay, which can detect multiple types of DNA damaging including single-strand and double-strand DNA breaks, loss of HORMAD1 led to a significant increase in tail moment following exposure to HU (Figure 4B). Together, these results indicate HORMAD1 protects against the generation of DNA breaks. We postulated that an increase in fork degradation and DSB formation would lead to the generation and/or exposure of ssDNA. Consistent with this notion, we observed that sgHORMAD1 cells exhibit elevated accumulation of ssDNA as measured by IdU intensity both at baseline and in response to HU (Figure 4C). To evaluate the consequences of this DNA damage on chromosomal integrity, we performed metaphase spreads. As expected of cancer cells, sgCTRL displayed a low but detectable incidence of chromosomal aberrations including fragmented chromosomes and chromatid breaks following HU treatment. However, accumulation of chromosomal aberrations was significantly increased both in untreated and HU exposed sgHORMAD1 cells. (Figure 4D). Collectively, our findings indicate that anomalously expressed HORMAD1 in LUAD cells protects DNA from damage due to replication stress that would otherwise lead to catastrophic chromosomal defects.

**Figure 4.**
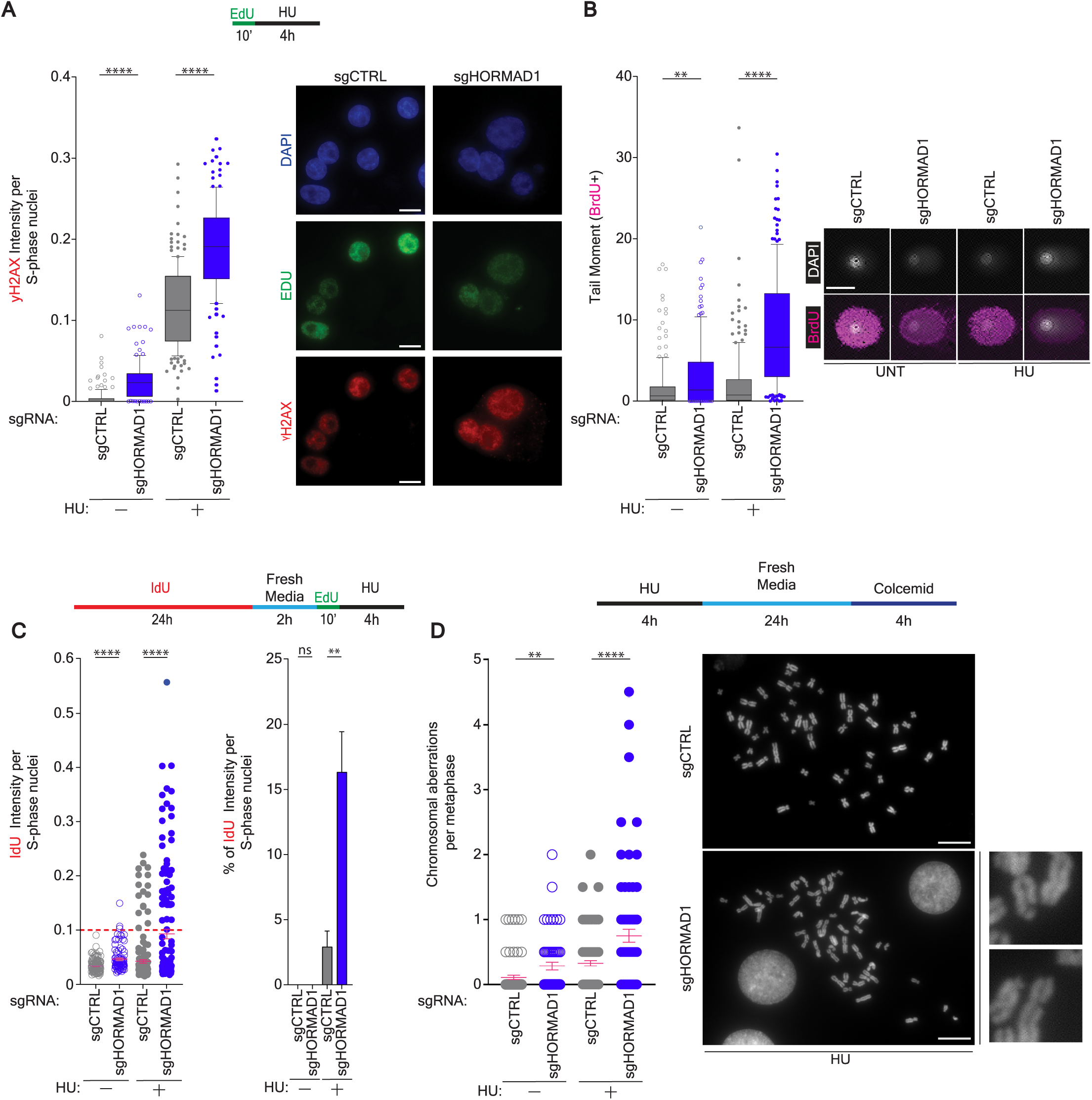
HORMAD1 mitigates DNA damage during DNA replication stress. (A) Indicated H1395 cells were labelled with EdU for 10 minutes then exposed to HU or fixed. Immunofluorescence was performed for mean γH2AX and EdU. Center line indicates the median, bounds of box indicate the first and third quartile, and whiskers indicate the 10th and 90th percentile for mean γH2AX nuclear fluorescence intensity in EdU positive cells. *P* value calculated by Mann-Whitney test; n≥3. (**B)** Indicated H1395 cells were labelled with BrdU for 20 minutes and fixed immediately or exposed to HU and fixed followed by an alkaline comet assay. Left: Box plot was derived from measuring the tail moment in BrdU positive cells and boundaries are indicated as in (A). *P* value calculated by Mann-Whitney test; n≥3. Right: Representative images of comets are shown. (Scale bar = 50μm) (**C)** Indicated H1395 cells were labelled as indicated in the diagram (top). Control cells were fixed following EdU labelling. Immunofluorescence was performed for IdU and EdU under non-denaturing conditions. Left: IdU nuclear fluorescence intensity in individual EdU positive cells. Mean is indicated. *P* value calculated by Mann-Whitney test; n≥3. Right: Mean IdU staining at or above a background threshold (0.1, red line in left panel) was plotted as a percentage of EdU positive cells. (**D**) H1395 cells were treated as indicated in schematic (top) prior to performing a metaphase spread assay. Each data point represents one metaphase spread. Mean is indicated. Error bars represent ± SEM. Right: Representative pictures at 100X with zoomed in examples. Scale bar = 10μm. *P* value calculated by Mann-Whitney test; n=3.

## DISCUSSION

Meiotic proteins are frequently aberrantly expressed in tumor cells. The contribution of these proteins to DNA replication and repair has only recently begun to be addressed (Lingg et al., 2022). Here, we report a novel function for the meiotic HR factor, HORMAD1, in the protection of stalled replication forks in cancer (Chen *et al.*, 2005b; Gao *et al.*, 2018; Liu *et al.*, 2020a; Nichols *et al.*, 2018; Tarantino *et al.*, 2022; Watkins *et al.*, 2015; Xian Wang, 2018; Zong *et al.*, 2021). This protection appears to be due to HORMAD1’s capacity to recruit BRCA2 and RAD51 to reversed forks. In developing gametes, HORMAD1 associates with DNA and serves as a platform for a multiprotein complex to recruit SPO11. In tumors, HORMAD1 could be enacting a similar protein complex assembly mechanism to stabilize forks against breaks or collapse. Moreover, it is possible HORMAD1 alters the required stoichiometry of the factors required for DNA fork protection. Indeed, the chronic state of replication stress in tumors has been implicated in limiting available pools of the protection factor, RPA (Toledo et al., 2017). Importantly, previous studies have implicated HORMAD1 in recruitment of RAD51 to γ-irradiation-induced DNA double strand breaks, suggesting a conserved mechanism in supporting RAD51 nucleofilament formation irrespective of the damage (Nichols *et al.*, 2018). Future studies will be necessary to elucidate the molecular mechanism by which HORMAD1 mediates RAD51 loading.

Another notable finding is the specificity for HORMAD1 function in fork protection. First, our data indicate that HORMAD1 acts specifically on forks reversed by the FBH1 helicase. Recent studies have indicated that fork reversal can be mediated by a number of different translocases or helicases in a non-redundant fashion (Liu *et al.*, 2020b). The substrate is likely dictating this specificity, which would suggest that HORMAD1 is specific to the FBH1-generated substrate. In this setting, HORMAD1 specifically protects against the MRE11 endonucleolytic mediated degradation coupled to DNA2 and BLM activity. As further details emerge regarding fork reversal mechanisms and the substrates that are generated, the basis for HORMAD1 selectivity could be illuminated.

We also observed a specificity among tumor-derived cell lines where HORMAD1 is required for stalled fork protection. Of the eight cell lines we tested, five exhibited a degradation phenotype upon HORMAD1 knockdown. Surprisingly, A549 cells, which require HORMAD1 for RAD51 loading following γ-irradiation were not dependent upon HORMAD1 for fork protection (Nichols *et al*., 2018). This phenotype extended to the molecular level as BRCA2 and RAD51 loading on stalled forks were not impacted in the absence of HORMAD1 (Figure S3). Collectively, our work suggests HORMAD1 may exhibit both substrate and context selectivity in its fork protection mechanism.

It is well established that HORMAD1 expression correlates with an increase in mutation burden, neo-antigen production, and gross chromosomal abnormalities in cancer (Gao *et al.*, 2018; Liu *et al.*, 2020a; Nichols *et al.*, 2018; Watkins *et al.*, 2015). One major question that has emerged from the compendium of studies is whether this correlation is cause or consequence HORMAD1 expression. Two reports have suggested HORMAD1 promotes genomic instability when inappropriately expressed by biasing cancer cells towards using the error-prone DNA repair pathways translesion synthesis (TLS) or non-homologous end joining (NHEJ), or by blocking DNA mismatch repair (Liu *et al.*, 2020a; Tarantino *et al.*, 2022; Xian Wang, 2018). Alternatively, evidence illustrates that HORMAD1 promotes HR-mediated repair of DSBs, suggesting it also contributes to genome maintenance (Gao *et al.*, 2018; Nichols *et al.*, 2018; Watkins *et al.*, 2015). Our study adds more evidence to the possibility that HORMAD1 promotes genome stability by reducing the accumulation of genomic errors due to replication stress. We reason that early in malignancy, cell growth is limited by excessive replication stress and ensuing DNA damage induced by oncogene activation (Kotsantis et al., 2018). Pre-malignant cells that engage HORMAD1 to protect stalled forks could gain a selective growth advantage. In support of this possibility, a recent study reported that chronic overexpression of cyclin E in normal, RPE, cells results in a period of slowed growth and replication stress, which is subsequently resolved. Transcriptomic analysis found that HORMAD1 expression significantly increased in cells that regain proliferative capacity despite cyclin E overexpression (Limas et al., 2022). This finding supports the notion that HORMAD1 expression is directly engaged to overcome a growth bottleneck associated with excessive DNA replication stress.

We postulate that HORMAD1’s function may be a mixture of genomic protection and disruption, depending on the stage and type of cancer or additional changes in DNA repair pathways (BRCA1/2 mutation, ATM/ATR loss, etc). The diversity of HORMAD1 function could be due to its capacity to bind to chromatin as well as a variety of replication and repair proteins and serve as a platform for protein complex formation. Despite the disparate findings in these studies, one theme is strikingly consistent: loss of HORMAD1 increases the sensitivity of tumors to DNA damage agents including IR, PARPi or 6-TG treatment, suggesting that targeting HORMAD1 could represent a mechanism to dramatically increase the log-kill of these anti-cancer agents without compromising their narrow therapeutic window(Gao *et al.*, 2018; Liu *et al.*, 2020a; Nichols *et al.*, 2018; Watkins *et al.*, 2015).

## MATERIAL AND METHODS

### Cell Lines and Culture conditions

All NSCLC cell lines were obtained from John Minna (UT Southwestern) between 2014 and 2018. NSCLC cell lines were maintained in Roswell Park Memorial Institute medium (RPMI-1640; Sigma Aldrich) supplemented with 5% FBS (Sigma Aldrich) and incubated at 37°C in a humidified 5% CO_2_ atmosphere. HEK293T cells were obtained from ATCC in 2014 and cultured in DMEM media supplemented with 10% FBS at 37°C in a humidified 5% CO_2_ atmosphere. All cells were periodically evaluated for mycoplasma contamination by a mycoplasma PCR detection kit (Cat# G238, ABM). Cells were genotyped at the beginning and end of the study.

### Inhibitors

Hydroxyurea (HU) (Chem Impex International (50-525-181)) was used at a final concentration of 4 mM for four hours. Olaparib (Thermo Fisher Scientific (NC0136853)) was used at a final concentration of 10 μM. The MRE11 Exonuclease inhibitor, Mirin, (Cayman Chemicals (13208)) was used at a final concentration of 50 μM. The MRE11 Endonuclease inhibitor, PFM01 (Sigma-Aldrich (SML1735-5MG)) was used at final a concentration of 10 μM. The DNA2 C5 inhibitor (AOBIOUS (NSC15765)) was used at a final concentration of 20 μM.

### Antibodies

Antibodies used: γH2AX (05-636, EMD Millipore) (Western Blot (WB) 1:1000, Immunofluorescence (IF) 1:1000), RPA2 (ab2175, abcam) (WB 1:1000, IF 1:500), RAD51 IF (ABE257, Sigma Aldrich) (1:100), RAD51 WB (D4B10, CST) (1:1000), HORMAD1 (HPA037850, Sigma-Aldrich) (1:5000), CHK1 (2360S, CST) (1:1000), pCHK1 (S345) (2348S,CST) (1:1000), ATR (2790S, CST) (1:1000), pATR (T1989) (58014S,CST) (1:1000), Histone 3 (4499S,CST) (1:1000), CldU (ab6326, abcam) (1:500), IdU (347580, BD Biosciences) (IF 1:50, DNA fiber assay 1:500), BrdU (5292S, CST) (1:100), Vinculin (sc-73614, Santa Cruz) (1:3000), GAPDH (G8795, Sigma Aldrich) (1:10000), BRCA2 (10741S, CST) (1:1000), MRE11 (NB100-142, Novus Biologicals) (1:1000), EXO1 (A302-640A, Bethyl Labs) (1:1000), DNA2 (ab96488, Abcam) (1:1000), BLM (A300-110A-M, Bethyl Labs) (1:1000), Beta-Tubulin (2128S, CST) (1:5000), Normal Rabbit IgG (2729S, CST), Ku80 was described previously (Lu et al., 2019).

### Transfections

For siRNA transfections, cells were trypsinized and seeded in Opti-MEM containing Lipofectamine RNAiMAX (Thermo Fisher Scientific, Waltham, MA) with a pool of siRNAs at a final concentration of 50 nM. siRNAs for were purchased from Sigma as follows: non-targeting controls (VC30002), targeting HORMAD1 (SASI_Hs01_00222714, SASI_Hs01_00222715,SASI_Hs01_00222716), MRE11 (SASI_Hs02_00339885), EXO1 (SASI_Hs01_00076269),DNA2(SASI_Hs02_00314950), WRN (SASI_Hs01_00219502), BLM (SASI_Hs01_00103851,SASI_Hs01_00103853), SMARCAL1 (SASI_Hs02_00328762, SASI_Hs02_00328763), HLTF (SASI_Hs01_00162286, SASI_Hs01_00162287), ZRANB3 (SASI_Hs01_00109807, SASI_Hs01_00109806, SASI_Hs01_00109804), FBH1 (SASI_Hs01_00057770, SASI_Hs02_00360372, SASI_Hs01_00057771, SASI_Hs01_00057772). If more than one siRNA is indicated, these were combined onto a pool and used for the respective experiment.

### Lentiviral Transduction

Stable cell lines were generated through lentiviral-mediated transduction. HEK293T cells were cotransfected with the target gene vectors (in pLX304) and lentiviral packaging plasmids (psPAX2 and pMD2.G). 48 hours later, virus-conditioned medium was harvested, passed through 0.45 μm pore-size filters, and then used to infect target cells in the presence of Sequa-brene (S2667, MilliporeSigma) for at least six hours. Six hours later, medium was replaced, and cells were allowed to recover. Stable sgCTRL and sgHORMAD1 cell lines were generated using pLX-sgRNA and pCW-Cas9 constructs (Addgene plasmid #50662, #50661) as described previously (Nichols *et al.*, 2018).

### Alkaline Comet assay

2 million cells were seeded on 10 cm^3^ dishes for 24 hours. Subsequently, 4 mM Hydroxyurea was added to the media for four hours prior to cell collection and comet assay in accordance with the manufacturer’s instructions (R&D #4250-050-K). Briefly, cells were suspended in 1.5 mL of cold PBS and diluted 1:10 dilution with low melting agarose. This mixture was placed on a slide and incubated at 4°C for 30 minutes. Slides were lysed overnight at 4°C with provided lysis buffer. Slides were incubated in alkaline unwinding solution at room temperature for 20 minutes (pH >12). Slides were then electrophoresed at 25 Volts for 30 minutes at 4°C in the dark. Slides were incubated in H2O twice for five minutes and 70% EtOH once. Slides were dried for 15 minutes at 37°C and then washed with two changes of PBS for 5 minutes each and blocked with blocking buffer (PBS containing 0.1% BSA and 0.1% Tween20), for 30 min at room temperature. The slides were incubated with mouse monoclonal anti-BrdU (1:100) diluted in blocking buffer in the dark in a humidified box at room temperature for 1 h. The excess of primary antibody was washed off with three changes of PBS and once with blocking buffer and probed with secondary antibody (Alexa Fluor 488-labelled goat anti-mouse antibody (1:1000) diluted in blocking buffer for 1 h. Following incubation, cells were washed as previously mentioned, air dried for 15 minutes at 37C, then mounted onto glass slides using ProLong Gold Antifade reagent with 4’,6-diamidino-2-phenylindole (DAPI). Images were acquired by epifluorescence microscope Keyence BZ-X700 at 10X magnification. Tail moments were scored using the CometScore software (TriTek)

### Immunofluorescence Assays

Immunoflourescence (IF) assays were used for visualizing γH2AX, RPA, and RAD51. Cells were plated on coverslips 24 hours before commencing the experiment to allow for attachment. The cells were treated with PBS or 4 mM of Hydroxyurea for four hours. For nuclear pre-extraction cells were washed with PBS and Cytoskeletal Buffer (CSK) (10 mM PIPES, pH 6.8, 100 mM NaCl, 300 mM sucrose, 3 mM MgCl_2_, +/− 0.3% Triton X-100) once. Cells were then incubated for five minutes with Triton-CSK buffer (0.3% Triton-X) and washed with CSK buffer (no triton) four times. Cells were washed in cold PBS and fixed at room temperature in 4% formaldehyde and blocked for 30 min at room temperature in blocking buffer (5% BSA in PBS). Primary antibodies were diluted with blocking buffer and incubated with samples overnight at 4°C. Following washes, samples were incubated with Alexa Fluor-conjugated secondary antibodies (Thermo Fisher Scientific) at a dilution of 1:1000 for one hour at room temperature. Where specified, 5-ethynyl-2’-deoxyuridine (EdU) (Cayman chemicals (#20518)) was used at final concentration of 20 μM for 10 minutes. Bromodeoxyuridine (BrdU) (Cayman chemicals (#15580)) was used at final concentration of 20μM for 20 minutes. Coverslips were then mounted onto glass slides using ProLong Gold Antifade reagent with DAPI. Images were acquired by epifluorescence microscope Keyence BZ-X700.

For parental ssDNA detection cells were seeded on coverslips in a 24-well plate for 24 hours. Subsequently, cells were pulsed for 24 hours with IdU (50 μM), washed with PBS, incubated in fresh media for two hours, and subsequently either incubated with 4 mM Hydroxyurea for four hours or immediately collected. Samples were pre-extracted and processed as described above for immunofluorescence in native conditions. Cells were then imaged using a Keyence BZ-X700 microscope at 60X magnification. Fluorescence intensity was measured using Cell Profiler Software.

### Cell Lysis and Western Blotting (WB)

Samples were lysed in preheated (100°C for five minutes) 2X Laemmli sample buffer with Beta-mercaptoethanol and boiled for five minutes. Samples were resolved using SDS-PAGE, and transferred to an Immobilon PVDF membrane (Millipore), blocked in 5% non-fat dry milk followed by incubation in the respective primary antibodies overnight. Following incubation, membranes were washed three times with Tris Buffered Saline (20 mM Tris, 150 mM NaCl, 0.1% Tween-20) (TBST), and incubated for one hour with horseradish peroxidase-coupled secondary antibodies (Jackson Immunoresearch). Subsequently, membranes were washed three times with TBST and then developed using SuperSignal™ West Pico PLUS chemiluminescence substrate (Thermo Fisher scientific, 45-000-875). Western blots were scanned using the EPSON perfection v700 photo scanner.

### DNA Fiber assays

Cells were pulse-labelled with 20 μM 5-iodo-2’-deoxyuridine (IdU) then 200 μM 5-chloro-2’-deoxyuridine (CldU) as indicated in the experimental schemes. For immunodetection of labelled tracks, the following primary antibodies were used: anti-CldU (rat monoclonal anti-BrdU/CldU; BU1/75 ICR1 Abcam, 1:500) and anti-IdU (mouse monoclonal anti-BrdU/IdU; clone b44 Becton Dickinson, 1:500). The secondary antibodies used were goat anti-rat Alexa Fluor 488 or goat anti-mouse Alexa Fluor 596 (Thermo Fisher scientific, 1:1000). The incubation with antibodies was accomplished in a humidified chamber for 1 hour at room temperature. ProLong Gold Antifade reagent with DAPI was used to mount slips on glass slides and images were acquired by a Leica DM5500 B upright microscope at a 60X magnification. Length of labelled tracks was measured using Image J, and a minimum of 100 individual fibers were analyzed per biological replicate. At least 3 biological replicates were performed.

### Cell Cycle analysis and Cell Number Analysis

Two million cells were seeded on a 10-cm^3^ dish 24 hours prior to the onset of the experiment. Cells were then mock treated or treated with 4 mM Hydroxyurea for four hours, and then washed once with PBS. Cells were then trypsinized and cell pellets obtained and washed again with PBS. Cells were fixed in ice-cold 70% EtOH at −20°C for 30 minutes. Cells were then washed with PBS and resuspended in DNA extraction buffer (0.2 M Na2HPO4, 0.1 M Citric Acid: pH 7.8) at room temperature for 20 minutes. Cells were subsequently centrifuged, supernatant was removed, and cells were stained with Propidium Iodide solution (Thermo Fisher Scientific (AAJ66584AB), 80 μg/ml; 0.1% Triton X-100; 100 μg/ml RNase A in PBS) in the dark for 20 minutes. Cells were immediately analyzed by flow cytometry using a BD LSR Fortessa instrument and BD FACSDiva 6.2 software. A minimum of 0.5 × 10^4^ cells were analyzed per condition. FlowJo software was used to generate flow charts and calculate KS-Max difference. For cell proliferation assays, cells were seeded at a density of 50,000 cells per well on a 6-well plate. Each day for six days a single well was trypsinized and counted to measure the number of cells. At least 3 biological replicates were performed. Cells were counted using Biorad TC-10 automated cell counter, and results were graphed using Graphpad prism.

### Subcellular Fractionation

Subcellular fractionations were performed using the Thermo-Fisher scientific subcellular fractionation kit (Cat#78840). Protein content was measured using the Pierce BCA protein assay kit (Cat#23225). SDS-PAGE and immunblotting. Blots were scanned on an EPSON perfection v700 photo scanner. Band intensity was measured using Image J for each respective protein band. Normalization was done first for loading using either Ku80 or Histone H3. p-ATR and p-CHK1 were normalized to their total unphosphorylated counterparts following normalization to total loading.

### Metaphase Spread

2 million cells were seeded in a 10 cm^3^ dish. 24 hours later, samples were treated with 4 mM Hydroxyurea or vehicle (PBS) for four hours, followed by a PBS wash and incubation in fresh media for 24 hours. Subsequently, cells were treated with 0.2 ug/ml Colcemid for 4 hours (KaryoMAX). Cells were then trypsinized and collected and incubated in prewarmed (37°C) 7.5 mM KCl for six minutes. Cells were resuspended in pre-chilled Carnoy fixative (3:1 methanol:acetic acid), pelleted and resuspended in cold Carnoys fixative. The samples were placed in a dropwise manner onto slides, allowed to air dry, and mounted in ProLong Gold Antifade reagent with DAPI. Metaphase spreads were then imaged using a Keyence BZ-X700 microscope at 100X magnification. Chromosomal aberrations of gaps, fragments, and breaks were quantitated manually.

### qPCR

RNA was extracted using GenElute Mammalian Total RNA Miniprep Kit (RTN350) (Sigma Aldrich) and reversed transcribed to cDNA using High-Capacity cDNA Reverse Transcription Kit (4368814) (Thermo scientific). Quantitative PCR was performed with triplicates in 96-well plate format on the QuantStudio 3 Real-Time PCR system (ThermoFisher Scientific). HORMAD1, SMARCAL1, HLTF, ZRANB3 and FBH1 quantitative PCR was performed using QuantiTect SYBR Green PCR kits. mRNA expression levels were calculated used the threshold cycle (2^−ΔΔCT^) method, normalized to RPL27.

### Liquid chromatography-tandem mass spectrometry (LC-MS/MS) based HORMAD1 interactome analysis

Cells were lysed for 30 min on ice in non-denaturing lysis buffer (NDLB) (50 mM HEPES pH 7.4, 150 mM NaCl, 1.0 % Triton X-100, 0.5% sodium deoxycholate, 25 mM β-glycerophosphate, 1 mM EDTA, 1 mM EGTA, 1 mM Na3VO4, 1 μg/mL pepstatin, 2μg/mL leupeptin, 2μg/mL aprotinin, and 10μM bestatin). Lysates were clarified at 12,000 x g for 10 minutes and pre-cleared with Protein A/G agarose beads for 1 hour at 4°C. Five percent of each lysate was set aside as input material and the remainder was immunoprecipitated with appropriate antibodies overnight at 4°C. Protein A/G agarose beads blocked with 0.5% BSA were added in the final two hours of immunoprecipitation. Beads were then washed three times in NDLB, followed by elution in 2X sample buffer containing β-mercaptoethanol. Immunoprecipitated proteins were separated by SDS-PAGE. After staining with Colloidal Coomassie blue, bands were excised from the acrylamide gel and in-gel tryptic digestion was performed. For mass spectrometry, samples were digested overnight with trypsin (Pierce) following reduction and alkylation with DTT and iodoacetamide (Sigma–Aldrich). The samples then underwent solid-phase extraction cleanup with an Oasis HLB plate (Waters) and the resulting samples were injected onto an Orbitrap Fusion Lumos mass spectrometer coupled to an Ultimate 3000 RSLC-Nano liquid chromatography system. Samples were injected onto a 75 um i.d., 75-cm long EasySpray column (Thermo) and eluted with a gradient from 0-28% buffer B over 90 minutes. Buffer A contained 2% (v/v) ACN and 0.1% formic acid in water, and buffer B contained 80% (v/v) ACN, 10% (v/v) trifluoroethanol, and 0.1% formic acid in water. The mass spectrometer operated in positive ion mode with a source voltage of 1.5 kV and an ion transfer tube temperature of 275 °C. Mass spectrometry (MS) scans were acquired at 120,000X resolution in the Orbitrap and up to 10 MS/MS spectra were obtained in the ion trap for each full spectrum acquired using higher-energy collisional dissociation (HCD) for ions with charges 2-7. Dynamic exclusion was set for 25 seconds after an ion was selected for fragmentation.

For H1395 cells, raw MS data files were analyzed using Proteome Discoverer v2.4 SP1 (Thermo), with peptide identification performed using Sequest HT searching against the human protein database from UniProt. Fragment and precursor tolerances of 10 ppm and 0.6 Daltons were specified, and three missed cleavages were allowed. Carbamidomethylation of Cys was set as a fixed modification, with oxidation of Met set as a variable modification. The false-discovery rate (FDR) cutoff was 1% for all peptides. For A549 cells, raw MS data files were converted to a peak list format and analyzed using the central proteomics facilities pipeline (CPFP), version 2.0.3(Trudgian and Mirzaei, 2012; Trudgian et al., 2010). Peptide identification was performed using the X!Tandem and open MS search algorithm (OMSSA) (Craig and Beavis, 2004; Geer et al., 2004)search engines against the appropriate protein database from Uniprot, with common contaminants and reversed decoy sequences appended (Elias and Gygi, 2007). Fragment and precursor tolerances of 20 ppm and 0.6 Da were specified, and three missed cleavages were allowed. Carbamidomethylation of Cys was set as a fixed modification and oxidation of Met was set as a variable modification. Label-free quantitation of proteins across samples was performed using SINQ normalized spectral index Software (Trudgian et al., 2011).

Peptides with an enrichment ≥ 1.4 (HORMAD1/IgG Control) and a minimum of 4% coverage were considered interactors.

### Statistical Analysis

Graphpad Prism (Graphpad Software) was used to perform statistical analyses. Outliers were removed by applying the outlier Shapiro-Wilk test from GraphPad prism. For Box and Whisker graphs, the center line indicates the median, bounds of box indicate the first and third quartile, and whiskers indicate the 10th and 90th percentile. Statistical differences in all cases were determined by Mann-Whitney, t-test, or a one-way Anova as specified in the figure legends.: ns= not significant, P > 0.05; *P < 0.05; **P < 0.01; ***P < 0.001; ****P < 0.0001.

## ACKNOWLEDGEMENTS

Mass Spectrometry was performed through the UTSW Proteomics Core. We would like to thank Andrew Lemhoff with help in the proteomic studies. Funding was provided by the National Institutes of Health to A.W.W. (CA196905; on which L.H.R. was supported with an administrative supplement) and to A.J.D. (CA162804 and GM04725) This work was also supported by The Welch Foundation (I-2087-20210327). Support for use of core services was made possible through NCI Cancer Center Support Grant (5P30CA142543) to the Harold C. Simmons Cancer Center.

---

**Supplemental Table 1.** List of HORMAD1 interacting partners involved in DNA metabolism in A549 and H1395 cells.

| DEAD Box Proteins | Description | References |
| --- | --- | --- |
| DDX50 | Responds to viral RNA | Han, P., et al. (2017) |
| DDX21 | Resolves R-loops, nucleolar | Song, C., et al. (2017) |
| DDX17* | Resolves DNA DSB by formation of DNA: RNA hybrids | Boleslavskaya, B., et al. (2022) |
| DDX3X*/DDX3Y | Resolves DNA: RNA hybrids; recruited to DNA damage sites | Bol, G. M., et al. (2015) |
| DDX5* | Resolves DNA: RNA Hybrids; BRCA2 interactor for DNA repair | Sessa, G., et al. (2021) |
| DDX41 | Prevents DNA damage, R-loop interactor, telomere maintenance | Mosler, T., et al. (2021) |
| DDX1 | Resolves DNA:RNA Hybrids, required for RPA, RAD51 loading | Li, L., et al. (2016) |
| **DNA Metabolism** |  |  |
| MCM3 | DNA helicase minichromosome maintenance complex | Meagher, M., et al. (2019) |
| MCM5 | DNA helicase minichromosome maintenance complex | Meagher, M., et al. (2019) |
| MCM6 | DNA helicase minichromosome maintenance complex | Meagher, M., et al. (2019) |
| MCM7 | DNA helicase minichromosome maintenance complex | Meagher, M., et al. (2019) |
| RUVBL1 | DNA helicase, DNA replication and radioresistance | Yenerall, P., et al. (2020) |
| PCNA* | Faithful duplication of DNA | Arbel, M., et al. (2021) |
| WRNIP1 | Stalled fork protection | Leuzzi, G., et al. (2016) |
| ATAD5 | PCNA regulation and genome integrity | Park, S. H., et al. (2019) |
| RFC3 | PCNA regulation and genome integrity | Arbel, M., et al. (2021) |
| RFC4 | PCNA regulation and genome integrity | Arbel, M., et al. (2021) |
| NPM1* | ribosome synthesis, genomic stability, cellular growth and stress response | Wang, R., et al. (2022) |
| XRCC5 | Involved in genome integrity | Fell, V. L. and C. Schild-Poulter (2015) |
| XRCC6 | Involved in genome integrity | Fell, V. L. and C. Schild-Poulter (2015) |
| PARP1 | Parylation enzyme involved in genome integrity | Ray Chaudhuri, A. and A. Nussenzweig (2017) |
| MATR3 | DNA and RNA Binding protein, SFPQ-NONO DNA damage signaling | Salton, M., et al. (2010) |
| SFPQ * | Genome integrity, SFPQ-NONO DNA damage signaling | Salton, M., et al. (2010) |
| NONO | Genome integrity, SFPQ-NONO DNA damage signaling | Salton, M., et al. (2010) |
| C1QBP | Genome integrity, stabilization and regulation of MRE11 | Bai, Y., et al. (2019) |
| **Ribonucleoproteins** |  |  |
| HNRNPD | DNA: RNA Hybrid resolution for DNA repair | Alfano, L., et al. (2019). |
| HNRNPA2B1 | Nuclear DNA sensor | Zhang, F., et al. (2020) |
| HNRNPA1 | RNA splicing, Metabolism & telomere length | Alfano, L., et al. (2019). |
| HNRNPM | RNA splicing & Metabolism | Ho, J. S., et al. (2021) |
| HNRNPC | RNA splicing & Metabolism | Fischl, H., et al. (2019) |
| HNRNPU | RNA splicing & Metabolism | Han, B. Y., et al. (2022) |
| **DNA Structure Unfolding (R-Loops, G-Quadruplexes)** |  |  |
| CNBP | Prevents G-quadruplex formation | Benhalevy, D., et al. (2017) |
| POLDIP3 | R - loop resolution | Bjorkman, A., et al. (2020) |
| TDRD3 | R - loop resolution | Yuan, W., et al. (2021) |

**Supplemental Figure 1.**
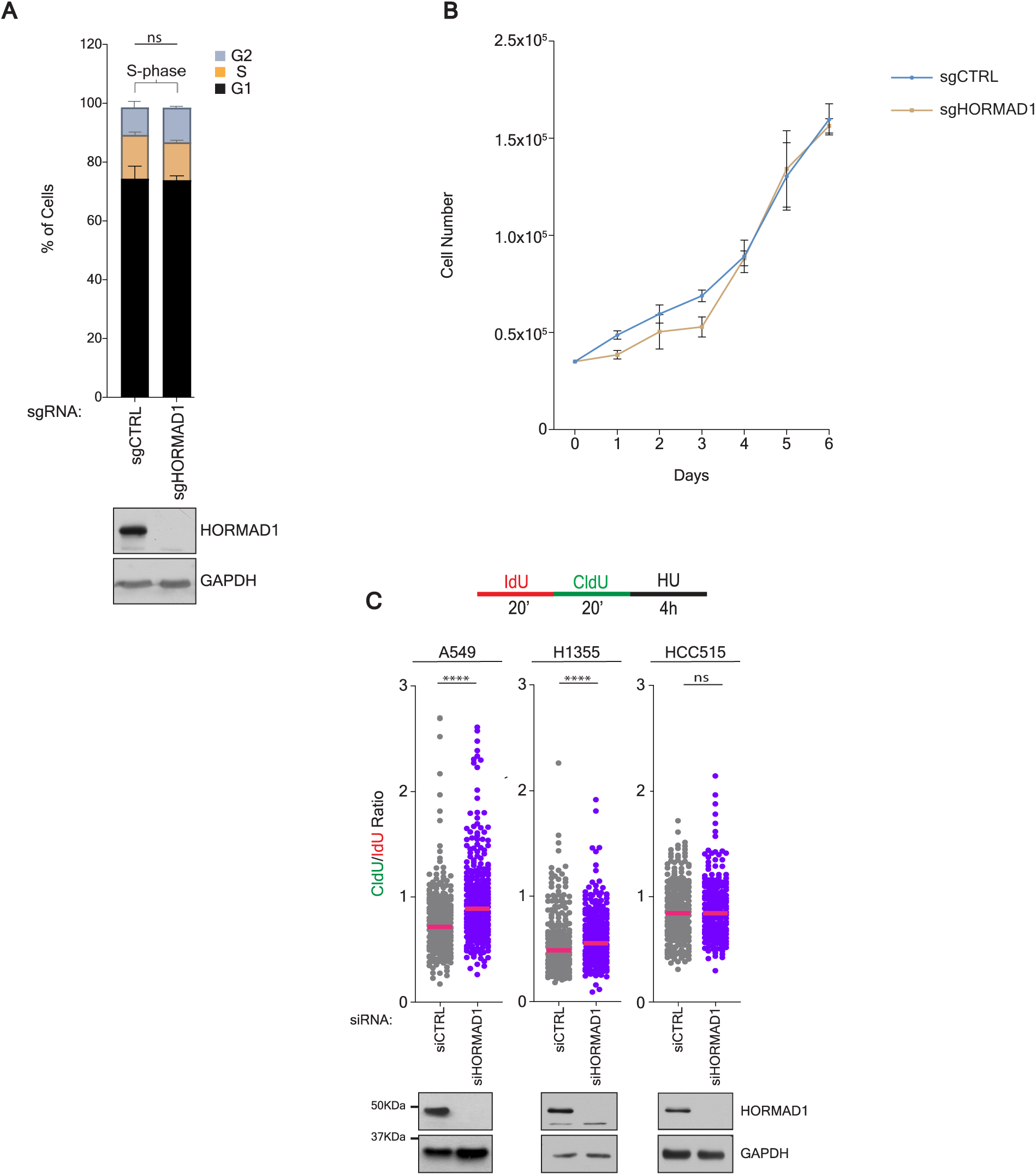
HORMAD1 knockout in H1395 lung adenocarcinoma cells does not affect cell cycle profile or cell proliferation. (A) Indicated H1395 cells were processed for cell cycle stage based in PI staining. Error bars represent SEM, n≥3. P value calculated by an unpaired t-test. (B) Indicated H1395 cells were seeded at density of 50k cells and counted each day for six days. Each point represents the mean cell number ± SEM for n≥3. P value calculated by an unpaired t-test. (C) A549, H1355, and HCC515 cells were transfected with indicated siRNAs for 72 hours, labelled as indicated in the schematic and a DNA fiber assay was performed. Each dot represents the ratio of CldU/IdU for a tract. Median indicated. P value calculated by Mann-Whitney t-test; n≥3. Below: Parallel lysates were obtained and western blotted for indicated proteins.

**Supplemental Figure 2.**
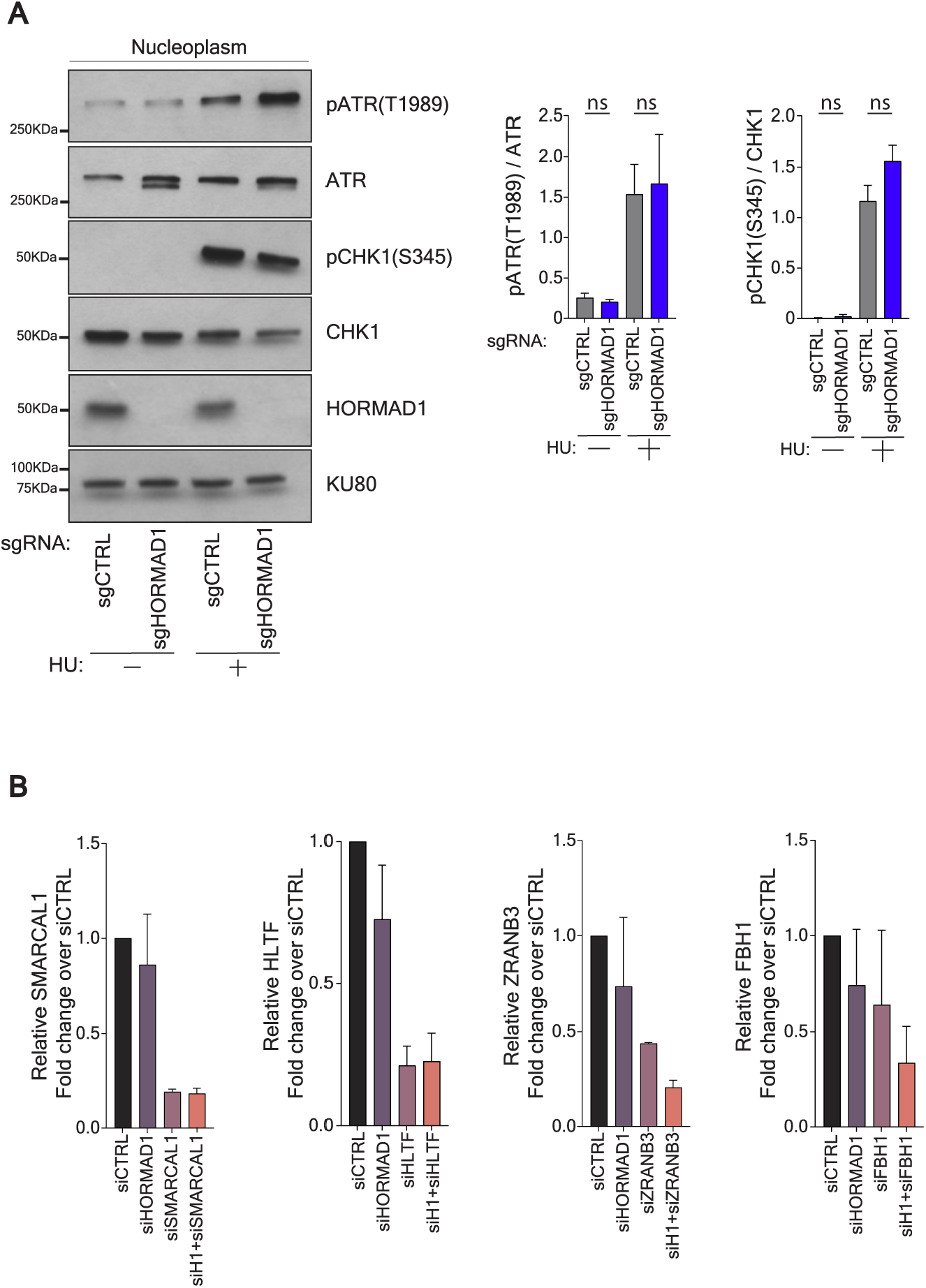
HORMAD1 acts downstream of checkpoint signaling and after FBH1 mediated fork reversal. (A) Indicated H1395 cells were exposed to PBS or HU for 4 hours, followed by subcellular fractionation. The nuclear fraction was subjected to immunoblotting with the indicated antibodies (Left). Graphs represent the mean band intensity over at least three experiments (Right). Error bars represent SEM. P value calculated by Mann-Whitney t-test; n≥3. (B) H1395 cells were transfected with indicated siRNAs for 72hours before a RT-PCR analysis was performed. The relative mRNA levels of HORMAD1, SMARCAL1, HLTF, ZRANB3 & FBH1 are shown for each indicated condition treated with indicated siRNAs, n=2. Mean is indicated & error bars represent range.

**Supplemental Figure 3.**
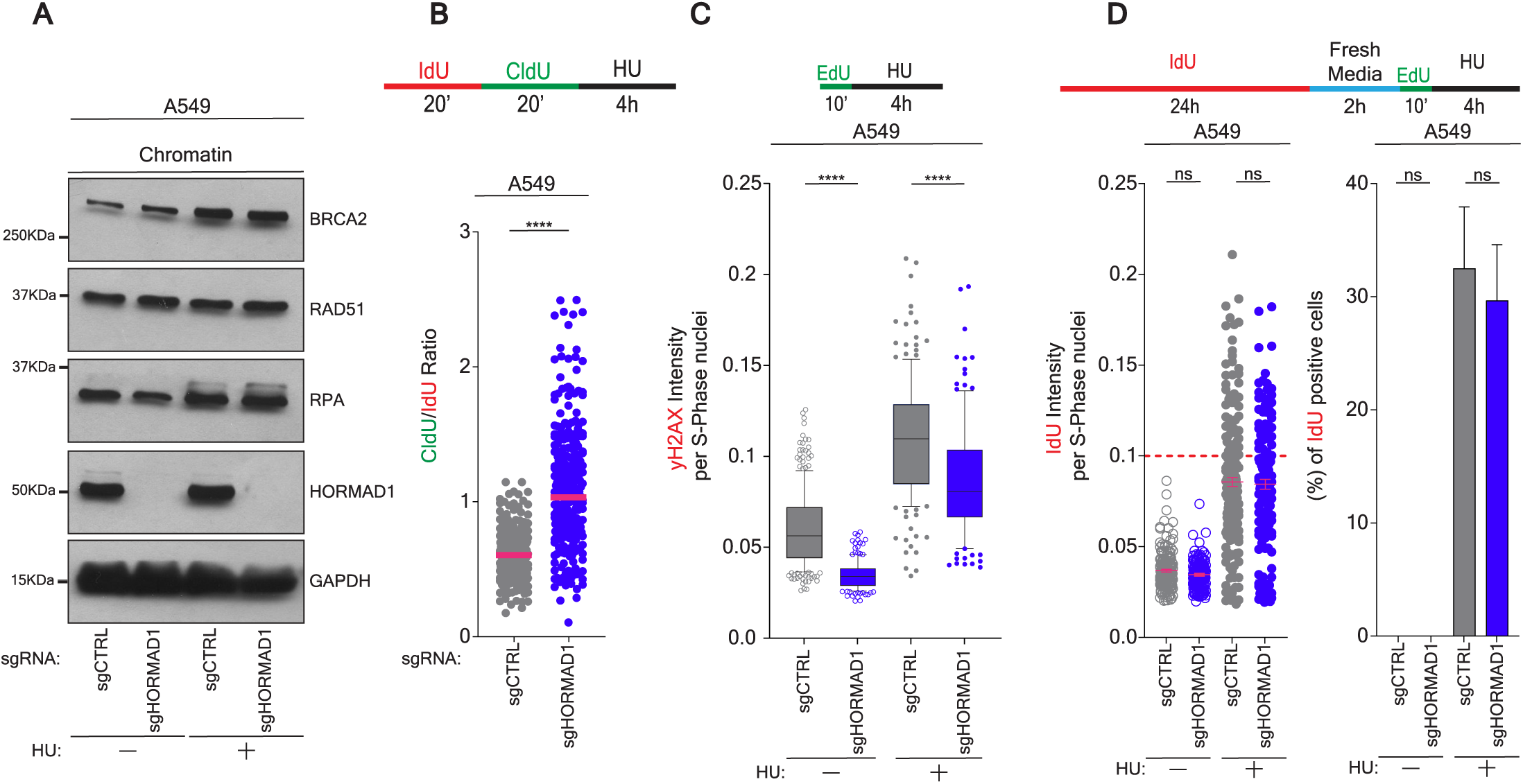
HORMAD1 is not required for BRCA2 or RAD51 chromatin loading or DNA integrity in A549 cells. Indicated A549 cells were exposed to PBS or HU for 4 hours, followed by subcellular fractionation. The nuclear fraction was subjected to immunoblotting with the indicated antibodies. Blots represent results from n=2. (B) A549 cells were treated as in the schematic (top). A DNA fiber assay was performed and CldU/IdU ratio graphed for each strand. P value calculated by Mann-Whitney t-test; n≥3. (C) Indicated A549 cells were exposed to PBS or HU for 4 hours prior to nuclear pre-extraction and fixation. Cells were exposed to EdU for 10 minutes prior to HU or collection. Immunofluorescence was performed forγH2AX and EdU. Box plot was derived from measuring the mean γH2AX nuclear fluorescence intensity in EdU positive cells. P value calculated by Mann-Whitney t-test; n≥3. (D) Indicated A549 cells were treated as in schematic (top) prior to nuclear pre-extraction and fixation. Immunofluorescence was performed for IdU and EdU under non-denaturing conditions. Mean IdU nuclear fluorescence intensity was measured in EdU positive cells (Left). Mean is indicated. Bar graph shows the mean number of IdU positive cells calculated as a percentage of all EdU positive cells at or above a minimum intensity (0.1 or dashed red line) (Right). Error bars represent SEM. P value calculated by Mann-Whitney t-test, n≥3.

